# Whole-Colony Modeling of Escherichia coli

**DOI:** 10.1101/2021.04.27.441666

**Authors:** Christopher J. Skalnik, Eran Agmon, Ryan K. Spangler, Lee Talman, Jerry H. Morrison, Shayn M. Peirce, Markus W. Covert

**Affiliations:** Department of Bioengineering, Stanford University, Stanford, CA 94305; Department of Bioengineering, Stanford University, Stanford, CA, United States; Department of Biomedical Engineering, University of Virginia, Charlottesville, VA, United States

## Abstract

Bacterial behavior is the outcome of both molecular mechanisms within each cell and interactions between cells in the context of their environment. Whereas whole-cell models simulate a single cell’s behavior using molecular mechanisms, agent-based models simulate many agents independently acting and interacting to generate complex collective phenomena. To synthesize agent-based and whole-cell modeling, we used a novel model integration software, called Vivarium, to construct an agent-based model of *E. coli* colonies where each agent is represented by a current source code snapshot from the *E. coli* Whole-Cell Modeling Project and interacts with other cells in a shared spatial environment. The result is the first “whole-colony” computational model that mechanistically links expression of individual proteins to a population-level phenotype. Simulated colonies exhibit heterogeneous effects on their environments, heterogeneous gene expression, and media-dependent growth. Extending the cellular model with mechanisms of antibiotic susceptibility and resistance, our model also suggested that variation in the expression level of the betalactamase AmpC, and not of the multi-drug efflux pump AcrAB-TolC, was the key mechanistic driver of survival in the presence of nitrocefin. We see this as a significant step forward in the creation of more comprehensive multi-scale models, and it broadens the range of phenomena that can be modeled in mechanistic terms.

**Author summary:** This work combines several models of molecular and physical processes that impact the physiology and behavior of the common microbe *Escherichia coli* into a multiscale model. Colonies comprised of multiple individual cells are simulated as they grow and divide—each with complex internal mechanisms, and with physical interactions and molecular diffusion in their environments. The integrative modeling methodology supports the addition of new submodels. The flexibility of this methodology is demonstrated by adding models of antibiotic resistance and simulating the colony’s response to antibiotic treatment.

## Introduction

Large-scale modeling of bacterial physiology has a history approaching 50 years, with Francis Crick first proposing to develop a complete “solution” of an organism such as *Escherichia coli* in 1973 [1]. Since that time, systems biology and computational modeling have grown in scope and scale to include integrative models of macromolecular assemblies [2], spatially-resolved stochastic models of cells [3], physics-based models of multicellular systems [4], genome-scale metabolic models [5], software for simulating whole cells [6], and models of minimal cells [7]. Wholecell models (WCMs), or integrative computational models with mechanistic and mathematical descriptions of entire cells [8], have built on these previous advances and raised the possibility of achieving Crick’s stated goal [9]. These models, which may be thought of as composites of diverse but highly integrated submodels that represent the known biological functions within a cell, are still very much works in progress [10]. That said, they have already succeeded in predicting novel experimental outcomes in *Mycoplasma genitalium* [8, 11] and *E. coli* [9]. Beyond prediction, whole-cell models also allow for the integration, reconciliation and even “deep curation” of massive, heterogeneous data sets—such as the decades of research performed using *E. coli* in labs around the world [9].

Previous whole-cell models have focused on the functions of single cells in isolation; however, many behaviors of *E. coli* can only be understood in the context of many bacteria interacting as a group. Examples include bet hedging [12], quorum sensing [13], “action potentials” in biofilms [14], and the response to antibiotics [15,16], to name a few. With regard to the latter, others have shown that stochastic expression of the multiple antibiotic resistance activator *marA* confers transient antibiotic resistance to single cells [17]. In a colony, heterogeneous *marA* expression has been shown both theoretically [18] and experimentally [17] to hedge against the possible appearance of antibiotics. Such phenomena can only be understood using a paradigm which accounts for cellular populations.

Agent-based modeling is an approach that simulates population behaviors that emerge from the actions and interactions of individual agents [19]. Agent-based models (ABMs) have been applied to blood vessel assembly [20], inflammation and fibrosis in human lungs [21], and human cancer tumors [22]. They have also been applied to bacterial populations, for example to study quorum sensing [23], chemotaxis [24], and colony growth [25].

Due to the complexity of the behaviors that are being simulated at the population level, the individual agents in ABMs typically follow a simple set of rules, although work by Peirce et al. has incorporated flux balance analysis (FBA) metabolic models with ABM [26]. What is required now is a method that builds upon this work by combining the full mechanistic detail of WCMs with ABM’s capacity to simulate emergent population behaviors. Here, we describe a new approach which allows us to synthesize ABM and WCM into a unified model. This approach was enabled by our novel model integration software, Vivarium [27]. We used Vivarium to construct a model of *E. coli* colonies where each cell was an instance of an *E. coli* model from the *E. coli* Whole-Cell Modeling Project [28] that interacted with other *E. coli* models in a shared spatial environment via physical forces and molecular exchange. We used the resulting “whole-colony” simulations to explore the impact of antibiotics on individual cells as well as on the colony as a whole.

## Results

We begin by describing our overall approach to simulating colony behavior, first in terms of the environment model and then in terms of modeling individual agents (for now, without antibiotic susceptibility or resistance). We then combine all of these models using the Vivarium framework, simulate them as an integrative multiscale model, and analyze the results. Finally, we add antibiotic susceptibility and resistance into the agent model’s functionality and run further simulations in order to test the model’s antibiotic response. Further details of our model, the simulations and the subsequent analysis can be found in the Methods section, as well as in the code itself, which is freely available at https://github.com/CovertLab/WholeCellEcoliRelease. [Note that the code is not yet available in the `WholeCellEcoliRelease` repository. It is currently available at https://github.com/CovertLab/wcEcoliColoniesTEMPORARY.]

### Simulating Colony Behavior

#### Modeling the Environment

Colony simulations contain capsule-shaped, bacterial agents that interact in a shared spatial environment via physical forces and molecular exchange (Fig 1). Colony growth is produced by the individual agents’ nutrient uptake, internal cellular functions, and cell division (Fig 1A). The environment and agents are two-dimensional for simplicity, but they each have a volume. In Vivarium, the simulation state is held in “state variables,” which are grouped into “stores.” The model is defined by “processes,” each of which updates various state variables at every time step. From here onward, we will use “process” to refer to a Vivarium process. The state variables for the environment and each agent are grouped into the stores shown in Fig 1B. Each agent is an instance of the same overall model, with states that evolve separately from any other agent. The independent evolution of these states allows heterogeneity to emerge across a growing colony as agents grow, divide and interact through their shared environment.

**Fig 1:**
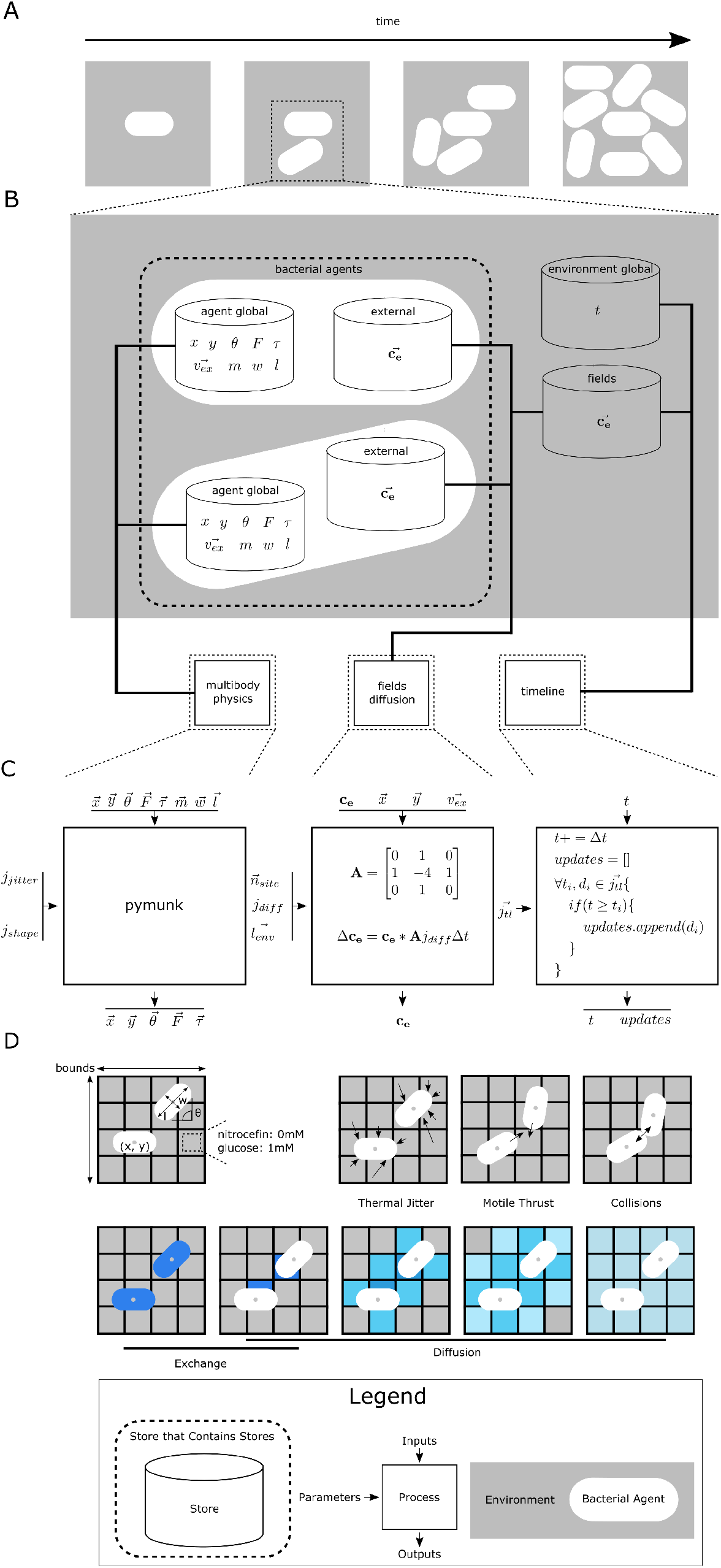
Environment model. Bacterial agents interact in a shared spatial environment by physical forces and molecular exchange. **(A)** Capsule-shaped agents interact and divide in a rectangular environment. **(B)** The environment model consists of a multibody physics process, a fields diffusion process, and a timeline process. These processes act on the global and external stores of each agent and on the environment’s global and fields stores. The dashed rectangle labeled “bacterial agents” represents a store that contains all the individual agents. **(C)** Specifications for environmental processes. The multibody physics process uses the pymunk physics engine to update each agent’s location and physical forces. The fields diffusion process models diffusion of molecules (**c**_e_) throughout the environment and exchanges 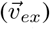 between the environment and agents. The timeline process updates the fields to model changes to the media condition according to a timeline 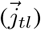. More detail is provided in the body text. **(D)** Cellular interactions mediated by the environmental model. The environment is a rectangular lattice (agents are drawn larger than each lattice site for ease of illustration, but in the actual model agents are smaller). Agent locations are continuous coordinates on this lattice, and agents have an angle from the x-axis, length, and width. Each lattice site has a concentration for each environmental molecule. Agents experience thermal jitter, motile thrust, and collision forces. Agents can also exchange molecules with the environment, and environmental molecules diffuse.

The model represents the environment as a two-dimensional rectangular space. Agent locations are continuous coordinates (*x, y*) within this space, and each agent has an angle from the x-axis (*θ*), length (*l*), width (*w*), thrust 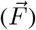, torque (*τ*), and mass (*m*). The environment is discretized into a lattice of sites, each with the same volume. The sites are larger than agents to reflect how even strong concentration gradients yield approximately equal concentrations over the small distances bacteria span [29]. Each site in the lattice has a concentration for each molecule in the environment.

Importantly, the agents do not interact directly; instead, all of their interactions are mediated by changes in the environment, where agents are buffeted by physical forces, molecules diffuse toward homogeneity, and the media can shift between environmental conditions (Fig 1C). The environment’s “multibody physics” process uses the pymunk physics engine, a two-dimensional rigid body simulator, to simulate physical interactions between cells [30]. We configured pymunk with force damping to simulate the low Reynolds number environment bacteria experience [31]; elasticity to simulate damped bacterial collisions; random jitter (*j_jitter_*) to model Brownian motion; capsule-shaped agents (*j_shape_*); and friction to model cell-cell adhesion. With these configurations, the multibody physics process updates the locations, angles, thrusts, and torques stored in each agent’s “globals” store, taking into account each agent’s mass, width, and length. The process is parameterized by the environment size 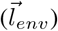, number of sites 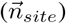, and diffusion rate (*j_diff_*). With regard to molecular diffusion, the “fields diffusion” process exchanges molecules between neighboring sites such that molecules move down concentration gradients. Molecule concentrations for each site in the environmental lattice are stored in the environment’s “fields” store. The process also updates each agent’s “external” store to reflect the concentrations in the cell’s local environment (site containing cell center), and exchanges molecules 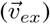 between agents (at locations 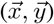) and the environment. Finally, the “timeline” process introduces external perturbations such as media shifts. The timeline process is parameterized by a time-stamped list of changes to make (*j_tl_*), and tracks the current time (*t*) in the environment’s “globals” store. The combined action of these processes enables agents to move through and exchange molecules with the environment, where the molecules diffuse (Fig 1D).

#### Modeling Bacterial Agents

Each bacterial agent is modeled by using a current source code snapshot from the *E. coli* Whole-Cell Modeling Project [28], which we extended by adding further processes in Vivarium (Fig 2). First, we discuss the processes needed to model colonies of bacteria, including one that contains the *E. coli* model. These core model processes (labeled with a “(1)”) appear in Fig 2, and the results of simulations that incorporate them are shown in Fig 3. The processes related to the antibiotic response (labeled with a “(2)”), are part of the extension described later when we discuss Fig 5. While the extension processes were part of the overall model that produced the results shown in Fig 3, they were not active since those simulations were antibiotic-free.

**Fig 2:**
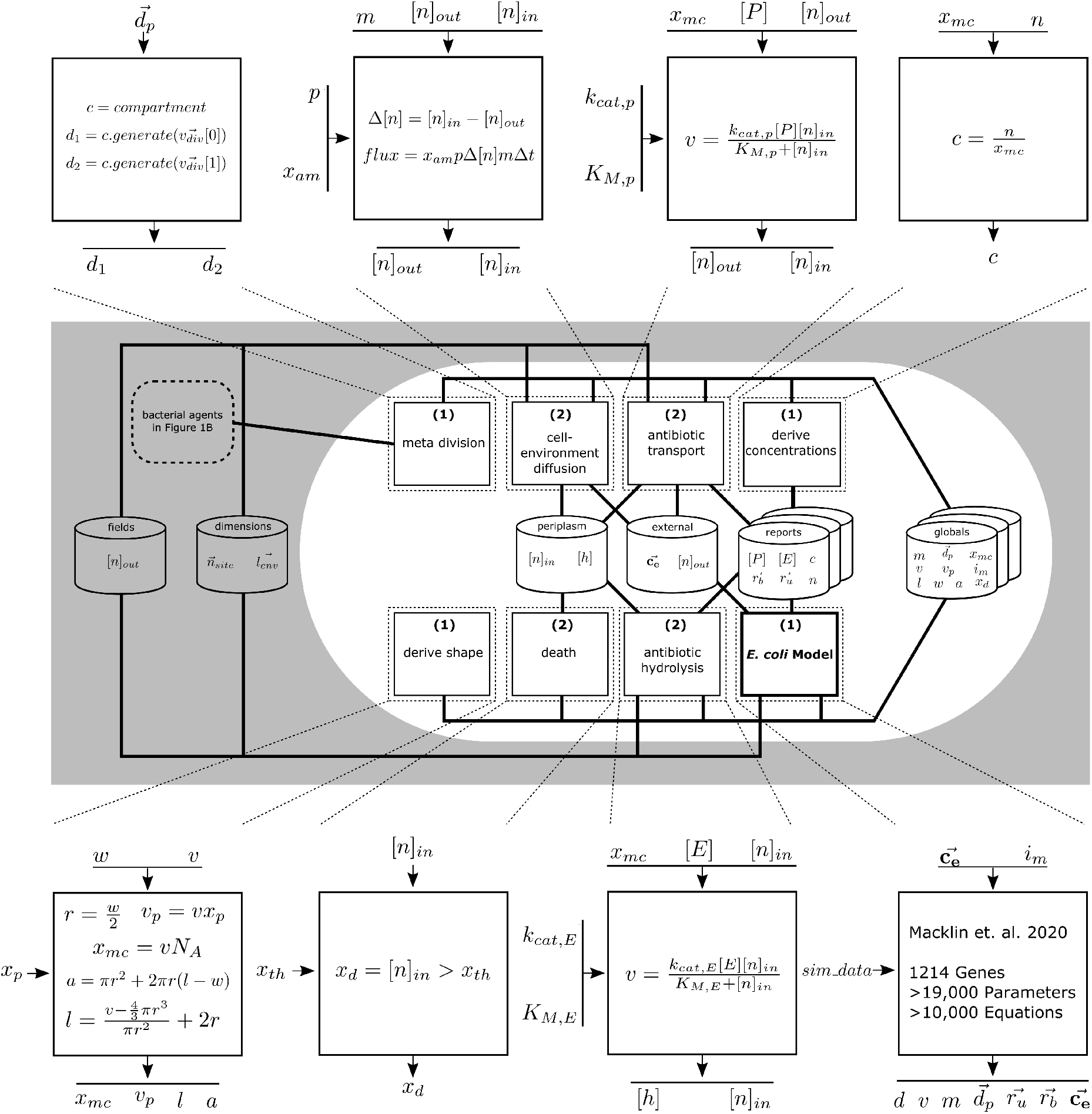
Bacterial agent model. Each bacterial agent is modeled by a set of core processes, plus extension processes to model antibiotic susceptibility and resistance. The core processes (labeled “(1)”) include: the *E. coli* process (bottom right), which represents internal molecular mechanisms using a snapshot from the *E. coli* Whole-Cell Modeling Project [28]; the derive shape process (bottom left), which calculates the cell’s length, surface area, a conversion ratio from concentrations to counts, and the periplasm volume; the derive concentrations process (upper right), which converts some of the counts in the reports store to concentrations; and the meta division process (upper left), which uses daughter cell states from the *E. coli* model to create agents for daughter cells. The extension processes (labeled “(2)”) include: the cell-environment diffusion process (upper middle-left), which uses Fick’s law to model antibiotic diffusing into the cell periplasm from the environment; the antibiotic transport process (upper middle-right), which models export of antibiotic from the periplasm to the environment by Michaelis-Menten kinetics; the antibiotic hydrolysis process (lower middle-right), which uses Michaelis-Menten kinetics to model antibiotic hydrolysis in the periplasm; and the death process (lower middle-left), which kills the cell once the internal antibiotic concentration exceeds a threshold. The equations in the figure describe how each process is implemented—these are covered in more detail in the body text.

**Fig 3:**
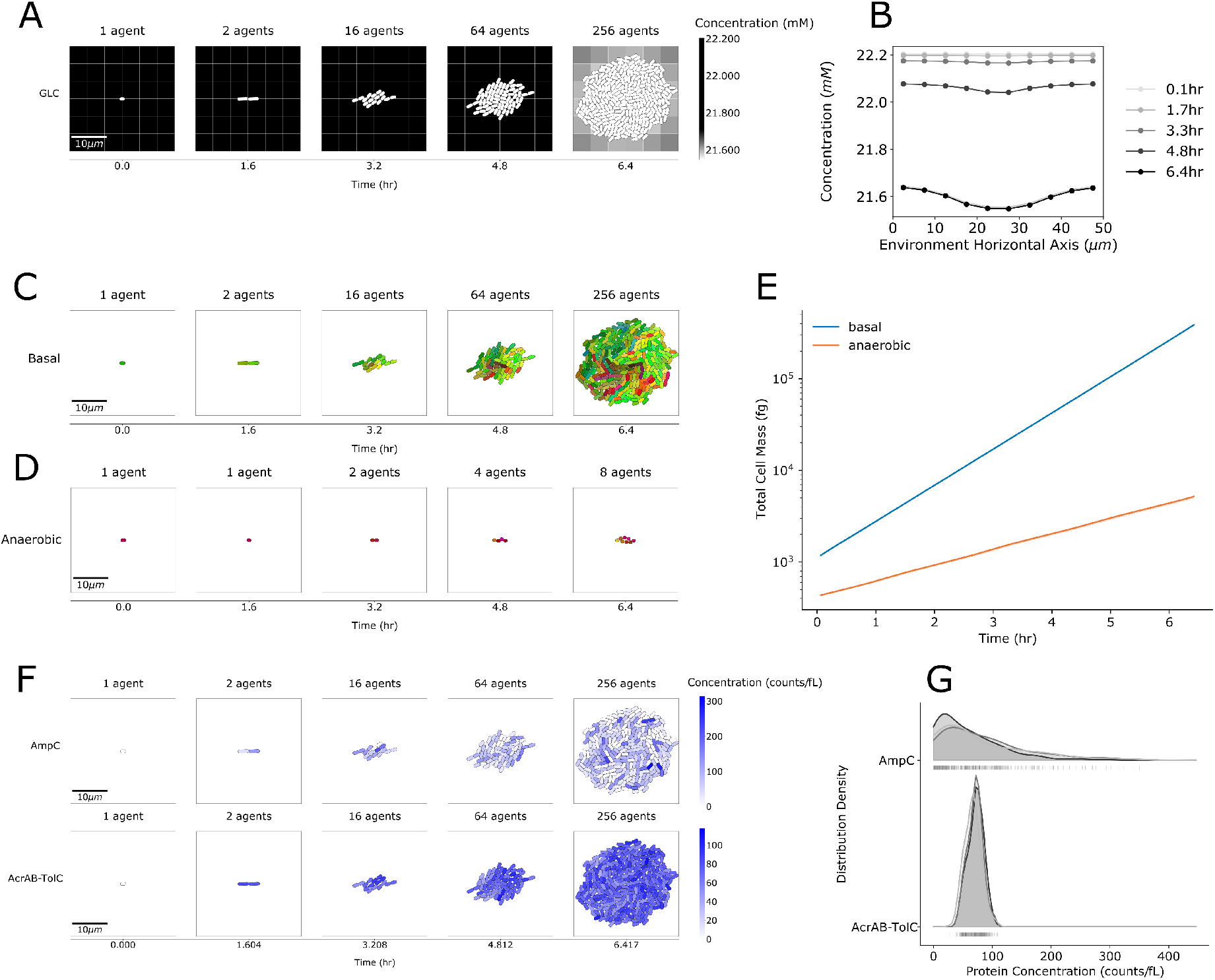
Heterogeneity in colony simulations. Panels (A), (C), (D), and (F) show representative examples from three simulations. **(A)** Snapshots of the colony over time. The cells are shown in white, and glucose concentrations are shown in grayscale for each lattice site. Bacterial agents have spatially heterogeneous effects on environmental molecules. **(B)** Consider a horizontal cross-section through the middle environment. For each of the snapshots in (A), the plot shows the glucose concentrations along the cross-section. The curve shows the median, and translucent bands show the IQR across the three simulations. The bands are too small to see without magnification. **(C)** Snapshots over time of the colony growing in basal conditions. Cell color represents phylogeny, with daughter cells inheriting their mother’s color with a slight offset in color space. **(D)** Snapshots over time of the colony growing in anaerobic conditions. **(E)** Total cell mass of the colony growing under basal and anaerobic conditions. The plot shows median colony masses with translucent bands for the IQR, though the bands are too narrow to see without magnification. **(F)** Snapshots of the colony over time with agents colored according to concentrations of the beta-lactamase AmpC and the multidrug efflux pump AcrAB-TolC. **(G)** Smoothed histograms of protein concentrations in the last snapshots of (F). Curves from each of the three simulations are overlaid in varying shades of gray. The vertical lines beneath the curves show the concentrations in each cell.

The current snapshot of the *E. coli* model (called “wcEcoli” here for convenience), built atop the one presented by Macklin et al. in 2020 [9], is based on 1214 gene functions and over 10,000 equations that describe the molecular mechanisms of *E. coli*. Since this version of wcEcoli was not created with Vivarium in mind, its internal state cannot be read by Vivarium processes directly. Thus, each *E. coli* process is a wrapper that holds an instance of wcEcoli. Every timestep, the *E. coli* process copies cell properties like mass (*m*), density (*d*), and some molecule counts (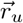 and 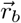) from wcEcoli to the store named “reports” that other processes can read. The *E. coli* process passes the concentrations 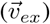 in the external store to wcEcoli, which uses them to calculate the amounts of molecules to exchange (import or export) with the environment. Imported molecules fuel metabolic reactions. The process uses the agent location (*x, y*), the environment’s size 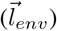, and the number of sites along each axis of the environment 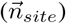 to apply the agent exchange molecules 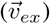 at the correct lattice site. wcEcoli is parameterized using `sim_data`, an object with model parameters calculated for wcEcoli. wcEcoli contains variant configurations for different environments; these environments are specified by a media identifier (*i_m_*) passed to wcEcoli as another parameter.

A process named “shape deriver” reads variables reported by the *E. coli* process, and it uses them to compute shape variables such as length and width. The *E. coli* process reports the cell’s volume (*v*), but other processes need more shape information. To calculate the cell’s surface area (*a*) and length (*l*), the shape deriver process imposes a capsule shape formed by a cylinder capped by hemispheres at each end. This process uses a constant width *w* = 1 *μ*m [32], so cells grow exclusively by elongation [33]. Shape deriver also calculates the concentrations-to-counts conversion ratio (*x_mc_*) by multiplying the cell volume by Avogadro’s number (*N_A_*) for other processes to use. Finally, the process uses a parameter *x_p_* = 0.3, the fraction of the cell’s volume consumed by the periplasm, to calculate the volume of the periplasm (*v_p_*) [34].

The “derive concentrations” process uses the concentration-to-counts conversion ratio (*x_mc_*) from the shape deriver to calculate concentrations. This is necessary because wcEcoli reports selected protein counts (*n*), but many of the processes expect concentrations (*c*).

Finally, to model cell division, the “meta-division” process instantiates a new agent for each daughter cell. When wcEcoli determines that the mother cell is ready to divide, it generates initial states for the two daughter cells, dividing the cell’s molecular contents between the two daughters using a binomial distribution to model stochastic partitioning [32]. This feature, combined with stochastic gene expression, facilitates the emergence of cellular heterogeneity as the colony grows. Once wcEcoli has saved the daughter cell initial states to files, it adds paths to these files to the 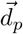 variable in the agent’s globals store. The “meta division” process uses the appearance of these paths as the trigger for cell division. The process creates a new agent from each daughter cell’s initial state and sets their locations such that the agents are end-to-end at the mother cell’s last position. It then returns these new agents (*d*_1_ and *d*_2_) to the Vivarium engine, which removes the mother cell agent from its environment and inserts the daughter cell agents.

### Simulations of Bacterial Agents Exhibit Environmental and Phenotypic Heterogeneity

With the completed Vivarium model, we first ran three 6.4-hour (8-generation) simulations in basal and anaerobic conditions. Since no antibiotics were present, the processes labeled “(2)” in Fig 2 were inactive. We found that our simulated colonies exhibited heterogeneous effects on their environments, heterogeneous expression, and media-dependent growth (Fig 3). In basal conditions (minimal media with glucose and oxygen), the colony grew to 256 cells (Fig 3A). Each agent had a doubling time of around 48 minutes.

Doubling time varied slightly because of stochasticity in wcEcoli. Since we did not model any causes of death (e.g. antibiotics) in this experiment, the colony grew to 2^8^ = 256 cells. We thought of the modeled system as a small colony growing on a large agar plate with plentiful nutrients, meaning that the constraints on nutrient uptake remained constant over the entire simulation. We achieved this by setting the volumes of lattice sites as if the environment were quite deep (1000 *μ*m), such that the glucose concentration changed very little (<1 mM) over the entire simulation.

As the colony grew, environmental glucose concentrations decreased as the sugar was consumed locally by the cells (Fig 3A,B). This decrease accelerated over time because of the colony’s exponential growth. Glucose concentrations decreased from an initial median of 22.2 mM (interquartile range, or IQR, across simulations: 1 × 10^−6^ mM) at 3.85 min to a final median of 21.6 mM (IQR across simulations: 9 × 10^−3^ mM) at 6.4 hr. Heterogeneity of glucose concentration increased over time from a median IQR of 6.85 × 10^−5^ mM (IQR across simulations: 7 × 10^−6^ mM) at time 3.85 min to a median IQR of 0.0343 mM (IQR across simulations: 2 × 10^−3^ mM) at 6.4 hr. While glucose was most strongly depleted from the center of the environment, concentrations on the periphery were also decreased by diffusion. This led to the formation of microgradients, which have also been observed experimentally [35].

When cells divide, their daughter cells are placed end-to-end. As they grow, each cell elongates and pushes upon its neighbors, which created a striated pattern of closely related cells proximate to each other (Fig 3C). However, thermal jitter disrupted this pattern, shifting individual cell alignment, and creating a round colony with imperfect striation. As can be seen in the figure, closely related cells still remained near each other, but with some mixing. This provided us with the ability to compare cells both by phylogenetic relation and spatial proximity—which comes into play in the later section on colony response to antibiotics.

We wanted to test the impact of the environment on colony size in our simulations. The current wcEcoli model supports three environments: glucose minimal with oxygen, glucose minimal without oxygen, and glucose media with amino acids and oxygen. Having performed our initial simulations in the first condition (basal, Fig 3C), we decided to observe the difference in colony growth in an environment without oxygen. In anaerobic media, the colony grew to only 8 cells over 6.4 hours (Fig 3D). At each time point for each condition, we computed the IQR of the colony mass across the three simulations and divided the IQR by the median colony mass. For each condition this value was smaller than 0.08, indicating little variation in colony mass across simulations. Colony growth in both conditions was exponential, but in anaerobic conditions the growth rate was much slower (Fig 3E). This observation matched our expectations and deepened our confidence in the simulations.

The protein concentrations in each agent are influenced by wcEcoli’s stochastic expression and partitioning (i.e., division) mechanisms, so protein expression was heterogeneous across the colony (Fig 3F). Such heterogeneity has also been observed experimentally [36]. As an example, we considered the case of the beta-lactamase AmpC and the multidrug efflux pump AcrAB-TolC, which protect the cell from beta-lactam antibiotics [37]. Certain proteins in this protective circuit have been shown to be expressed heterogeneously, both experimentally [17] and computationally [18]. Similarly, we found that AmpC also exhibited heterogeneity in expression across bacterial agents, with interquartile ranges between 68 and 100 proteins/fL in all three simulations (Fig 3G). The distribution of AcrAB-TolC concentrations was more consistent across simulations, with an IQR of 15-20 proteins/fL in each simulation. Importantly, the concentration of AmpC dropped to zero in 1.3% of the simulated cells; this never occurred for AcrAB-TolC.

### The Antibiotic Susceptibility and Resistance Model

We incorporated susceptibility and resistance to antibiotics into our bacterial model, following a conceptual framework in which cells import, export, and hydrolyze antibiotics (Fig 4) [37]. Briefly, antibiotics are allowed to diffuse into each agent cell’s periplasm through porins in the cell wall. Cells resist antibiotics using beta-lactamases that hydrolyze beta-lactams like nitrocefin [38] and multidrug efflux pumps that export antibiotics into the environment [39]. Hydrolyzed antibiotic is considered in our model to be inert (i.e., neither detrimental nor beneficial to the cell), while exported antibiotic returns to the environment, where it can diffuse back into cells. The periplasmic antibiotic concentration is thus increased by import and decreased by hydrolysis and export. If the periplasmic concentration reaches a parameterized threshold, the cell dies immediately. While dead cells remain in the simulation, they no longer grow, and their intracellular functions cease. These simplifications ignore the biological realities that many proteins, including beta-lactamases, can also function extracellularly [40] and that many other antibiotic resistance mechanisms operate in parallel [37]. We parameterized our model for nitrocefin, but this approach is meant to be applicable to other beta-lactams as well.

**Fig 4:**
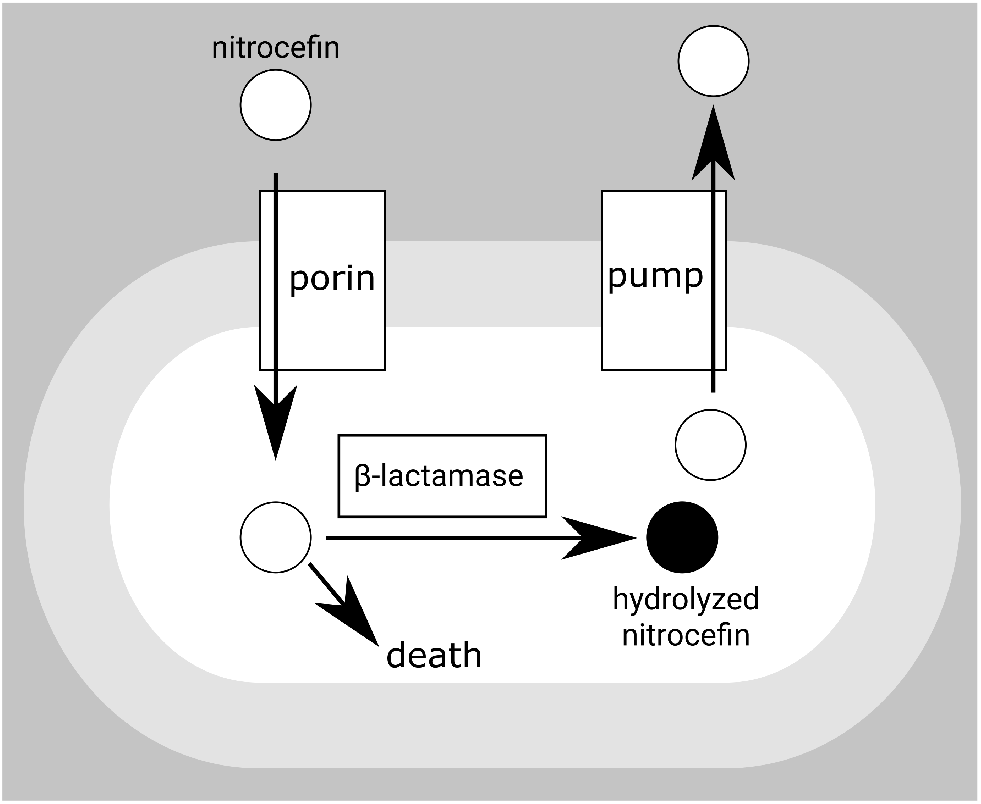
Model of antibiotic susceptibility and resistance. Nitrocefin diffuses into the periplasm through porins, is exported by the AcrAB-TolC pump, and is hydrolyzed by the AmpC beta-lactamase. If sufficient unhydrolyzed nitrocefin accumulates in the periplasm, the cell dies.

### Model Processes Related to the Antibiotic Response

The import, export, hydrolysis, and death mechanisms from Fig 4 are modeled by processes in each agent. These processes are labeled “(2)” in Fig 2.

The “cell-environment diffusion” process models nitrocefin import as Fickian diffusion from the environment into cells. Fick’s law uses the membrane’s permeability *p* = 2 × 10^−7^ cm/sec [41], which abstracts over the counts of the many porins in *E. coli* [42], and the cell’s surface area, which the process calculates from the cell mass (*m*) using the conversion ratio *x_am_* = 132 cm^2^/mg [41]. The concentration difference Δ[*n*] is calculated from the nitrocefin internal ([*n*]_*in*_) and external ([*n*]_*out*_) concentrations. Then by Fick’s law, the molecular flux is *v* = *mx_am_p*Δ[*n*]Δ*t* for timestep Δ*t*. The cell-environment diffusion process updates the internal and external nitrocefin concentrations to apply this flux.

The action of multidrug efflux pumps is modeled by the “antibiotic transport” process and follows Michaelis-Menten kinetics [43]. We parameterized the Michaelis-Menten equation with a rate constant *k_cat,p_* = 10 s^−1^ and a Michaelis constant *K_M,p_* = 4.95 × 10^−3^ mM [41]. The Michaelis-Menten calculations are performed using convenience kinetics [44]. The process uses the internal concentration of nitrocefin ([*n*]_*in*_) as the substrate concentration and the concentration of the export pump AcrAB-TolC ([*P*]) as the enzyme concentration. Updates to the external nitrocefin concentration ([*n*]_*out*_) are performed as exchanges, so the process uses the concentration-to-counts conversion ratio (*x_mc_*) and the timestep to convert the reaction rate (*v*) to a flux of nitrocefin out of the cell.

The action of the beta-lactamase is modeled by the “antibiotic hydrolysis” process as a Michaelis-Menten system. We parameterized the process with a rate constant *k_cat,E_* = 4 × 10^2^ s^−1^ and a Michaelis constant *K_M,E_* = 5 × 10^2^ *μ*M [45]. The process uses the internal concentration of nitrocefin ([*n*]_*in*_) as the substrate concentration and the concentration of the beta lactamase AmpC ([*E*]) as the enzyme concentration. Hydrolyzed nitrocefin is not an input to any process, which renders it inert.

We adopted a simple model of death where a cell dies instantly once the concentration of unhydrolyzed antibiotic in its periplasm ([*n*]_*in*_) exceeds a tolerance threshold (*x_th_*). After death, the cell body remains in the environment and continues to occupy space, but the *E. coli* model, meta division, death, antibiotic transport, and hydrolysis processes are removed to model the halting of the cell functions these processes implement. Processes are removed using Vivarium’s `_delete` operation, which removes processes but does not affect any stores or variables.

### Heterogeneity in Protein Expression Drives Cell Sensitivity to Antibiotics

We used the antibiotic model to test the susceptibility of our *in-silico* colonies to antibiotics. For the first half of each simulation, the colony grew without nitrocefin. Then, 3.2 hours into the simulation, nitrocefin was applied uniformly over the environment at the minimum inhibitory concentration (MIC), 0.1239 mM [46]. The only parameter that was left free in our model was the tolerance threshold (*x_th_*). To estimate this parameter value, we used a parameter scan including five tolerance thresholds ranging from 0.01 mM to 0.05 mM, with three simulations for each threshold.

The simulation results showed that when cells had lower tolerance for internal nitrocefin, colony growth was more sharply reduced (Fig 5A). With a tolerance threshold of 0.01 mM, all agents died after the antibiotic appeared and reached a median final mass of 21 pg (IQR across simulations: 0.9 pg), while at a threshold of 0.05 mM, colony growth slowed only slightly and reached a median final mass of 271 pg (IQR across simulations: 53 pg). Therefore, to reproduce a MIC of 0.1239 mM, the scan predicted that the tolerance threshold must be between 0.01 mM and 0.05 mM.

**Fig 5:**
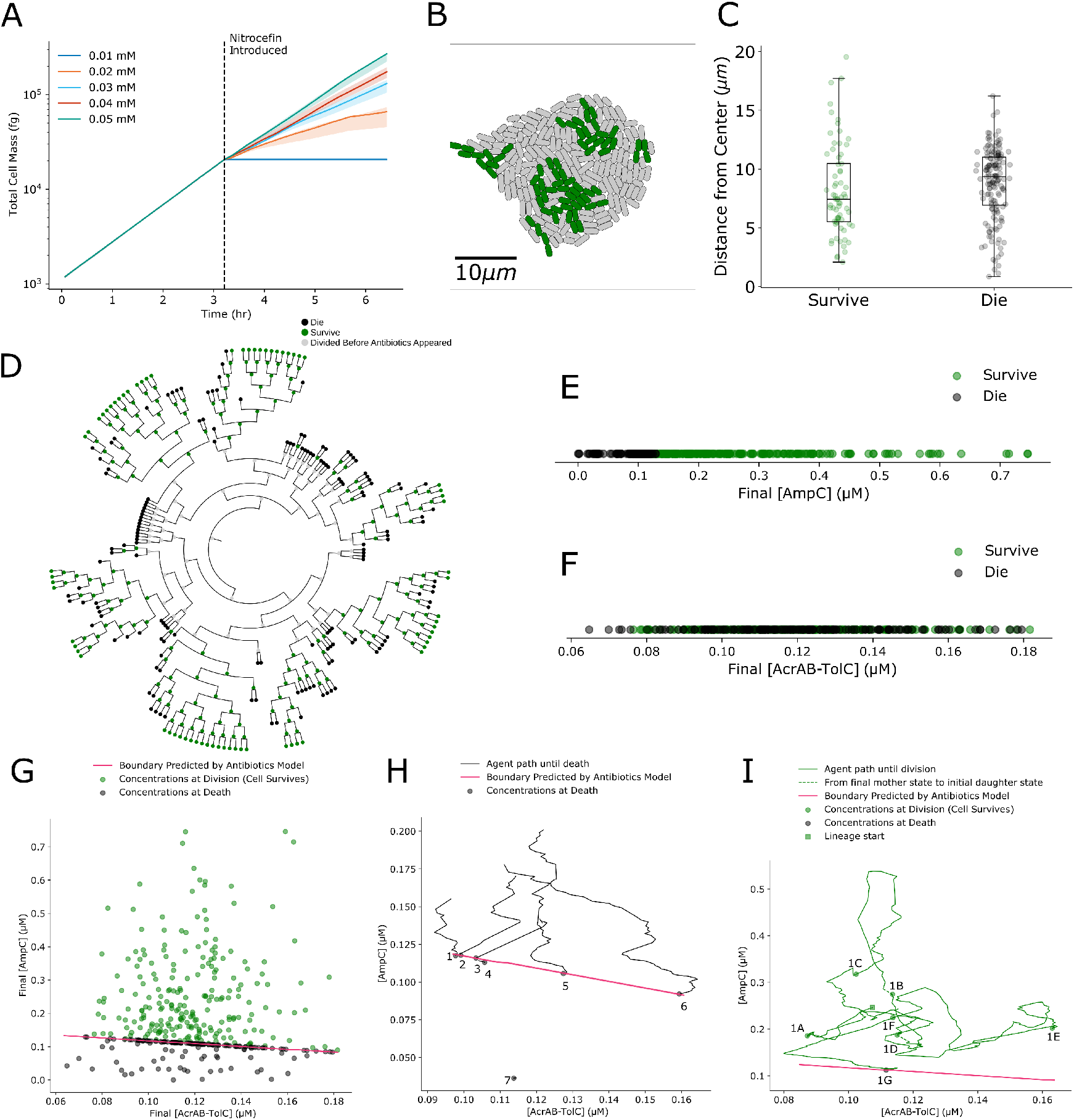
**(A)** Parameter scan for the lethal threshold of periplasmic nitrocefin. Each curve is a set of three simulations, each set run at the given threshold value shown by the line color. The curves indicate the median of the simulations, with translucent bands showing the IQR. Panels B-I are from a longer simulation with a threshold of 0.02 mM and plot cells that survived in green and cells that died in black. **(B)** The colony at the end of a 10.2 hour simulation, with living cells in green and dead cells in gray. Clustering of living cells could be a spatial effect or a phylogenetic effect. **(C)** Box plots of the radial distances between agents and the environment center. **(D)** Phylogenetic tree of all cells in the simulation. Cells that die from antibiotics are in black, those that survive are in green, and those simulated before the introduction of antibiotics are in gray. **(E)** The final AmpC concentrations for each cell that experienced antibiotics is shown as a dot. **(F)** The final AcrAB-TolC concentrations for each cell that experienced antibiotics is shown as a dot. **(G)** Plot of final AmpC concentrations against final AcrAB-TolC concentrations for each cell. The magenta curve shows the boundary between concentrations that agents could and could not survive, which we get from our parameterization of the antibiotic model in Fig 4. **(H)** Trajectories showing the concentrations experienced over time by agents that died. The dots indicate the agents’ final concentrations. Note that while time is not actually shown on the plot, distance on the plot approximates time since concentrations are changed continuously by expression events and dilution. The trace of agent 7 begins at a concentration where it cannot survive. It was at this concentration when nitrocefin was applied, or it could have been placed into that space by cell division. **(I)** The concentrations experienced by a lineage of cells over their lifetimes, beginning at the square with cell 1A and its progeny leading to cell 1G. Dots show each cell’s final concentrations, the solid lines show each cell’s history, and dashed lines connect mother and daughter cells.

These parameter scan simulations also exhibited substantial variation in colony mass. As in Fig 3E above, at each time point for each threshold concentration, we calculated the IQR of the colony mass and divided the IQR by the median colony mass. The maximum such value for each tolerance threshold was 0.05 for 0.01 mM, 0.4 for 0.02 mM, 0.3 for 0.03 mM, 0.3 for 0.04 mM, and 0.2 for 0.05 mM. Particularly for a threshold of 0.02 mM, these are much larger than we saw without antibiotics in Fig 3E. Thus, incorporation of the antibiotic model introduced variability into the simulation outputs—notwithstanding that the antibiotic extension processes are in fact deterministic.

We were intrigued by this variability and wanted to better understand how some cells lived while others died in our simulations. We ran a longer simulation (10.2 hours) with an antibiotic concentration of 0.1239 mM and a tolerance threshold of 0.02 mM. By the end of the simulation, 68.8% of the cells in the colony were dead (Fig 5B). Dead cells could be observed throughout the colony, with surviving cells forming clusters (because surviving cells continued to grow and divide over time). At the end of the simulation, nitrocefin concentrations had changed very little (over a range of 9 nM overall).

We next looked for features of individual cells that could be used to predict cell death. First, we considered each cell’s spatial location within the colony. The Euclidean distances of cells from the environment center showed medians of 7.4 *μ*m (IQR: 4.9 *μ*m) for live cells and 9.3 *μ*m (IQR: 4.1 *μ*m) for dead cells (Fig 5C). A Mann-Whitney U-test between these two groups yields a p-value of 0.039. This test had a power of 0.51 to detect a difference of 0.5 between the probability of death in the center and the periphery of the colony with α = 0.05. This is evidence of a possible correlation linking antibiotic resistance to spatial location within the colony.

We next considered the effects of cell lineage and ancestry. A phylogenetic tree analysis showed that some clusters of closely related cells survived until the end of the simulation, while other clusters of closely related cells were more likely to die (Fig 5D). Treating whether cells died as a binary trait, we calculated a phylogenetic signal of K = 1.35 (*p* ≤ 1 × 10^−5^) [47]. Therefore, whether a cell dies is highly correlated with its phylogenetic history.

We then sought to determine the mechanism of this correlation, focusing in particular on expression of the major proteins of interest, AmpC and AcrAB-TolC. We found that cells with a final (i.e. at death or division) AmpC concentration below 0.13 *μ*M tended to die, while cells with more AmpC tended to survive long enough to divide (Mann-Whitney U-test: *p* < 1 × 10^−54^, power > 0.99 for α = 0.05). In contrast, cell death was uncorrelated with final AcrAB-TolC concentration (Fig 5F). The distributions of final AcrAB-TolC concentrations in cells that survived until division and cells that died are indistinguishable (Mann-Whitney U-test: p = 0.65, power > 0.99 for α = 0.05). These results suggest that AmpC played a major role in determining which agents resisted nitrocefin, while AcrAB-TolC played little role.

To further explore the interactions between AmpC and AcrAB-TolC concentration in determining cellular antibiotic sensitivity, we derived a theoretical boundary between life and death (Fig 5G-I, magenta line) by analyzing a simplified model of the antibiotic response (which lacked the *E. coli* model) and assuming a tolerance threshold of 0.02 mM. We confirmed that cells with concentrations of AmpC and AcrAB-TolC that remained above this boundary throughout the simulation survived, while those with concentrations below the boundary died (Fig 5G). Moreover, when we traced the individual cell histories of dead cells through the space of AcrAB-TolC and AmpC concentrations (Fig 5H), we found that they died soon after crossing the boundary (Agents 1-6). We saw that the boundary was close to parallel with the AcrAB-TolC axis, confirming that the AcrAB-TolC concentration had little effect on whether a cell survived and that the AmpC concentration was the dominant factor.

Finally, we also considered the individual histories for surviving cells (Fig 5I). These trajectories form a quasi-random walk through the space of concentrations, with short jumps between generations due to stochastic partitioning upon division (dashed lines, for example between final concentrations in Agent 1B and initial conditions of progeny Agent 1C). Again, we see that these cells can live under virtually any of the cellular AcrAB-TolC concentrations, but die soon after crossing the concentration boundary for AmpC.

## Discussion

In summary, we integrated whole-cell and agent-based modeling to build models of *E. coli* colonies grounded in molecular mechanisms. We then extended this model with mechanisms of antibiotic susceptibility and resistance. The result was the first “whole-colony” computational model that mechanistically links expression of individual proteins to a population-level phenotype. The core colony model includes stochastic expression from the *E. coli* model (wcEcoli), and when we extended this core model with deterministic antibiotic processes, we observed an increase in the variation across trials. This occurred because the antibiotic processes depend on values such as AcrAB-TolC concentrations set by wcEcoli, and give function to the model’s stochasticity. Our simulations showed the growth of a colony in the presence of antibiotics, in which cells that did not express sufficient resistance proteins were eliminated from the population by death, and those that maintained higher resistance could continue to grow and divide. We see this as a significant step forward in the creation of more comprehensive multi-scale models of whole cells and their environments [48].

Our model also suggested that variation in the expression level of beta-lactamase AmpC, and not of multi-drug efflux pump AcrAB-TolC, was the key mechanistic driver of survival in the presence of nitrocefin. This result agrees with the finding by Nagano et al. that at the internal concentrations lethal to cells, efflux is negligible [41], but it stands opposed to other findings that efflux is much more important than hydrolysis in antibiotic resistance [49]. However, our finding comes with several caveats. Most notably, our model predicted a maximum survivable internal nitrocefin concentration (threshold) between one and two orders of magnitude higher than reported in the literature. Nagano et al. reported a threshold of 0.5 *μ*M for nitrocefin [41], which is consistent with the thresholds of 1.7 *μ*M for ampicillin and 0.36 *μ*M for benzylpenicillin reported by Kojima et al [49]. In contrast, our model predicted a threshold between 10 and 50 *μ*M (Fig 5A). This range arose from the interaction between the modeled mechanisms and other measured parameters, including the permeability of the cell wall to nitrocefin [41], the kinetic parameters for AmpC [45] and AcrAB-TolC [43], and parameters for their mRNA and protein expression and degradation in wcEcoli [9]. The discrepancy arises either due to the reported parameters being inaccurate or the modeled mechanisms being incomplete.

An extended model could include some of the additional mechanisms by which *E. coli* resist antibiotics, which would reduce the tolerance threshold needed to reproduce the minimum inhibitory concentration, bringing the threshold towards reported values. Three candidate mechanisms for future work include regulatory adaptation to antibiotics, the formation of protective biofilms, and evolution. First, *E. coli* respond to beta-lactam antibiotics through regulatory adaptation by inducing the expression of MarA, which activates downstream antibiotic resistance genes such as *micF*. MicF represses expression of the porin OmpF, but induces expression of the efflux pump AcrAB-TolC [50]—leading to less diffusion of antibiotics into the cell and more export out. An existing mathematical model of this mechanism [51] could be integrated in our model to capture this effect. Second, *E. coli* biofilms have been shown to protect against antibiotics through limited antibiotic penetration, reduced cell growth in nutrient-depleted microenvironments, and persister subpopulations which can tolerate high doses of antibiotics [52]. An important step toward incorporating these behaviors into our model would be making growth rate control mechanisms more responsive. Third, colonies can evolve antibiotic resistance or acquire it through horizontal gene transfer. El Meouche et al. found that cells with higher *acrAB-tolC* expression had lower expression of *mutS*, which is important for DNA mismatch repair [53]. This suggests that when exposed to antibiotics, *E. coli* have an increased mutation rate and thus evolve resistance quickly. We could extend the model with this mechanism by mutating upon division the basal transcription rates that parameterize the *E. coli* process.

By synthesizing whole-cell modeling and agent-based modeling using Vivarium, this work integrated mechanisms and parameters at the scale of protein and metabolic networks with mechanisms at the scale of small colonies. Future work should expand this integration both downward, to include molecular-scale models of protein structure and dynamics, and upward, to the scale of larger populations with multiple microbial species and complex microenvironments. A large body of work in structural biology has already demonstrated the utility of integrative approaches applied to structural models of macromolecular complexes [2]. This could extend the present model by connecting structural properties to constrain the behavior of reactions in the *E. coli* model. At the scale of populations, frameworks such as COMETS 2 [54] demonstrate how to model multiple microbial species in spatially structured environments by integrating dynamic flux balance analysis, molecular diffusion, and extracellular enzyme activity. We hope that the ongoing integration of models at these different scales will bring us continually closer to Crick’s dream of a “complete solution”.

## Methods

### Model

The model is fully defined by the code in the `WholeCellEcoliRelease` repository on GitHub (https://github.com/CovertLab/WholeCellEcoliRelease). Results can be reproduced using version `v2.0-beta.2` in that repository (DOI: 10.5281/zenodo.4695018. [Note that this code has not yet been released to the `WholeCellEcoliRelease` repository. It’s currently available at https://github.com/CovertLab/wcEcoliColoniesTEMPORARY/releases/tag/v2.0-beta.2.] The colony/compartments/antibiotics.py file defines the agent model, and the `colony/compartments/antibiotics_simple.py` file defines a simplified agent with the antibiotics model but without the full *E. coli* model. `The colony/experiments/antibiotics.py` file defines the environment model. It can also be executed to run a simulation. Its command-line arguments are documented in the file.

### Simulations

All simulations were run by executing the `colony/experiments/antibiotics.py` file with various command-line arguments. Simulations ran on a Google Compute Engine virtual machine (VM) with 96 CPUs and 442 GB memory. The instance boot disk had a capacity of 2000 GB and used a debian-10-buster-v20200618 image. The instance was also configured with a 500 GB swap file. Simulation data were emitted to a MongoDB server running on another Google Compute Engine VM. The MongoDB VM was of type g1-small (1 vCPU, 1.7 GB memory) and had a 500 GB boot disk with the image debian-9-stretch-v20191210.

The simulations we ran are described in Table 1, and each was executed using the following command:

~~~
OPENBLAS_NUM_THREADS=1 python -m \
colony.experiments.antibiotics -s SECONDS -o IP \
--pulse_concentration PULSE --update_fields \
[--anaerobic] --seed SEED --antibiotic_threshold \
THRESHOLD --emission EMISSIONS
~~~

**Table 1:** Parameters for all simulations. The “Time points” column lists the number of times data was collected over a simulation. The “Duration” column lists the total simulation time in seconds. The “[Antibiotic]” column lists the concentration, in mM, of antibiotic applied half-way through the simulation. The “Maximum survivable [antibiotic] threshold” column lists the tolerance threshold parameter set for agents in the simulation. The “Media” column shows the type of media the colony grew on. The “Seed” column shows the random seed used. The “Simulation” identifier. column shows a string that uniquely identifies the simulation.

| Time points | Duration (s) | [Antibiotic] (mM) | Maximum survivable [antibiotic] threshold (mM) | Media | Seed | Simulation identifier |
| --- | --- | --- | --- | --- | --- | --- |
| 100 | 23100 | 0 | n/a | basal | 0 | 20201119.150828 |
| 100 | 23100 | 0 | n/a | basal | 1 | 20210112.185210 |
| 100 | 23100 | 0 | n/a | basal | 2 | 20210125.152527 |
| 100 | 23100 | 0 | n/a | anaerobic | 0 | 20201221.194828 |
| 100 | 23100 | 0 | n/a | anaerobic | 1 | 20210205.163802 |
| 100 | 23100 | 0 | n/a | anaerobic | 2 | 20210205.183800 |
| 100 | 23100 | 0.1239 | 0.01 | basal | 0 | 20201228.172246 |
| 100 | 23100 | 0.1239 | 0.01 | basal | 1 | 20210209.174715 |
| 100 | 23100 | 0.1239 | 0.01 | basal | 2 | 20210209.192621 |
| 100 | 23100 | 0.1239 | 0.02 | basal | 0 | 20201228.211700 |
| 100 | 23100 | 0.1239 | 0.02 | basal | 1 | 20210210.181817 |
| 100 | 23100 | 0.1239 | 0.02 | basal | 2 | 20210210.210112 |
| 100 | 23100 | 0.1239 | 0.03 | basal | 0 | 20201229.160649 |
| 100 | 23100 | 0.1239 | 0.03 | basal | 1 | 20210212.160943 |
| 100 | 23100 | 0.1239 | 0.03 | basal | 2 | 20210212.193824 |
| 100 | 23100 | 0.1239 | 0.04 | basal | 0 | 20201230.191552 |
| 100 | 23100 | 0.1239 | 0.04 | basal | 1 | 20210217.152358 |
| 100 | 23100 | 0.1239 | 0.04 | basal | 2 | 20210217.201244 |
| 100 | 23100 | 0.1239 | 0.05 | basal | 0 | 20210206.045834 |
| 100 | 23100 | 0.1239 | 0.05 | basal | 1 | 20210219.151929 |
| 100 | 23100 | 0.1239 | 0.05 | basal | 2 | 20210220.153658 |
| 1000 | 36812 | 0.1239 | 0.02 | basal | 0 | 20210329.155953 |

`SECONDS` was the simulation duration (in seconds), `IP` was the internet protocol (IP) address of the MongoDB server, `PULSE` was the concentration of antibiotics (in mM) to apply, `--update_fields` specified that the exchange from the *E. coli* model should be applied to the environment, `--anaerobic` specified that the media should be anaerobic instead of basal, `SEED` was the random seed, `THRESHOLD` was the periplasmic antibiotic concentration cells can survive (in mM), and `EMISSION` was the number of times during the simulation to emit data for later analysis (i.e. the number of time points). When no antibiotic was applied (`PULSE` was 0), the value of `THRESHOLD` was irrelevant, so the `--antibiotic_threshold` argument was omitted.

### Analyses

As an alternative to running all the simulations, the raw simulation data are available as an archive under DOI 10.5281/zenodo.4697519. This archive identifies experiments by the identifier in Table 1 and can be passed to the analysis code described below. This archive was created as follows for a MongoDB server at IP address `IP`:

~~~
python -m src.archive_experiments -o IP
~~~

We analyzed properties of the simplified antibiotics model using the `colony/experiments/antibiotics_boundary_search.py` file in the `WholeCellEcoliRelease` repository. For particular AcrAB-TolC concentrations, this script searched for the AmpC concentration above which cells would survive and below which cells would die (assuming a threshold of 0.02 mM and an antibiotic pulse of 0.1239 mM). We executed this script on the analysis machine described below, and we will refer to the output file as `search.json`. This script was executed as follows:

~~~
python -m colony.experiments.antibiotics_boundary_search \ search.json
~~~

All of the figures and statistics in this paper were generated using version `1.0-beta.2` of the code in the `wcecoli-colony-analysis` repository on GitHub (https://github.com/CovertLab/wcecoli-colony-analysis/releases/tag/v1.0-beta.2). [Note that for calculating statistics, we applied commit `6ab4d5d417a03053758ff12c96dc090eb2320868` on top of version `1.0-beta.2` to fix a bug. We will soon include this fix in an updated release.] This version of the code has DOI 10.5281/zenodo.4706184.

The `src/make_figures.py` script contains the simulation identifiers in Table 1, and it uses these to retrieve simulation data from a MongoDB database or simulation data archive. We configured the GitHub Actions continuous integration service to run our analyses based on the archived simulation data. The `src/make_figures.py` script was executed as follows:

~~~
xvfb-run python -m src.make_figures SEARCH \
--data_path DATA --all
~~~

`DATA` was the path to the simulation data files and `SEARCH` was the path to `search.json`. The `xvfb-run` command is part of the X virtual framebuffer, which was required for the phylogeny plots. This script wrote the figures and a JSON file (`stats.json`) with statistics to a folder `out/figs`. We will refer to this path as `OUT`. Summary statistics were computed by executing:

~~~
python -m src.analyze_stats OUT/stats.json \
-o OUT/summary_stats.json
~~~

This wrote the summary statistics to `OUT/summary_stats.json`, which was human-readable. Lastly, phylogenetic signal was computed using the `src/analyze_phylogeny.r` like this:

~~~
RScript src/analyze_phylogeny.r OUT/phylogeny.nw \
OUT/agent_survival.csv
~~~

This script used `phytools` [47] and printed its output to the console. We used R version 4.0.4 and phytools version 0.7-70.

The `src/analyze_stats.py` script performed statistical tests using Scipy and Numpy [55, 56]. It also calculated the power of Mann-Whitney U-tests. First, the power of the test comparing Euclidean distances of live and dead cells from the environment center was calculated using the bootstrap method. We generated uniformly random cell locations within a 10 *μ*m radius of the origin. Let *l* be the number of live cells and *d* be the number of dead cells at the end of the simulation (Fig 2B). For each cell location, we randomly assigned it to be alive or dead with a probability of death that increased linearly from 0.25 in the center to 0.75 at a radial distance of 10 *μ*m from the origin. We then performed a U-test to compare the radial distances from the origin of the first *l* live cells versus the first d dead cells. We repeated this procedure 10,000 times, and our power was the fraction of these U-tests with a p-value below α. Second, the power of the test comparing concentrations of AcrAB-TolC and AmpC between live and dead cells was calculated using bootstrap as well. We used the average IQRs from the expression distributions in Fig 3D to estimate the standard deviations of AcrAB-TolC and AmpC concentrations, assuming a normal distribution. We used these standard deviations to create two normal distributions, one for each protein, with means separated by 0.1 *μ*M. We sampled from these distributions 10,000 times and estimated the power as the fraction of these samples where the Mann-Whitney U-test p-value was below α.

## Supporting information

**Table 2: S1 Table.** Variables shown in Figure 1.

| Variable | Description |
| --- | --- |
| $(x, y)$ | An agent’s location in the environment. |
| $\theta$ | Angle between an agent’s long axis and the environment’s x-axis. |
| $\vec{F}$ | The thrust force acting on an agent. |
| $\tau$ | The torque acting on an agent. |
| $v_{ex}$ | Molecular exchange, in counts. |
| $m$ | Agent mass. |
| $(w, l)$ | Agent size as its width and length. |
| $\vec{c}_e$ | Vector of environmental concentrations matrices. Each concentration matrix is a two-dimensional matrix with one cell for each site of the environment. |
| $t$ | Current simulation time, which is updated in discrete steps by the timeline process. |

**Table 3: S2 Table.** Parameters for the processes in Figure 1.

| Parameter | Value | Description |
| --- | --- | --- |
| $j_{jitter}$ | $1 \times 10^{-18}\ \text{N}$ | Force applied at random positions around each agent to model Brownian motion. |
| $j_{shape}$ | segment | A string describing the shape, which will represent each agent’s perimeter. |
| $j_{diff}$ | $0.5\ \mu\text{m}^2/\text{s}$ | Diffusion constant. |
| $\vec{j}_{tl}$ | varies | List of timeline elements. Each element consists of a time $t_i$ and an update $d_i$. |
| $n_{site}$ | $(10, 10)\ \mu\text{m}$ | Vector in $\mathbb{R}^2$ with the number of sites along the x and y axes in the environment. |
| $\vec{l}_{env}$ | $(50, 50)\ \mu\text{m}$ | Vector in $\mathbb{R}^2$ with the lengths of the x and y axes of the environment. |

**Table 4: S3 Table.** Variables shown in Figure 2.

| Variable | Description |
| --- | --- |
| $\vec{d}_p$ | A vector of 2 paths to daughter cell configurations produced by wcEcoli. |
| $d_1, d_2$ | Instantiated daughter cell agents that Vivarium can use to perform cell division. |
| $m$ | Agent mass. |
| $[n]_{out}$ | Concentration of nitrocefin outside the cell. |
| $[n]_{in}$ | Concentration of nitrocefin inside the cell. |
| $x_{mc}$ | The ratio of the count of a molecule to the molecule's concentration in the cell. |
| $[P]$ | Concentration of the AcrAB-TolC efflux pump. |
| $n$ | A molecule's count. |
| $c$ | A molecule's concentration. |
| $(w, l)$ | Agent size as its width and length. |
| $v$ | Total agent volume. |
| $v_p$ | Volume of the agent's periplasm. |
| $a$ | Agent surface area. |
| $x_d$ | Boolean describing whether the agent is dead. |
| $[E]$ | Concentration of the AmpC beta-lactamase. |
| $[h]$ | Concentration of the hydrolyzed nitrocefin. |
| $\vec{c}_e$ | Vector of environmental concentrations matrices. Each concentration matrix is a two-dimensional matrix with one cell for each site of the environment. |
| $i_m$ | One of a pre-specified set of media identifiers that specifies the media in which the agent is growing. |
| $d$ | The cell's density. |
| $\vec{r}_u$ | Reports of the counts of various molecules represented as unique molecules in wcEcoli. |
| $\vec{r}_b$ | Reports of the counts of various molecules represented as bulk molecules in wcEcoli. |

**Table 5: S4 Table.** Parameters for the processes in Figure 2.

| Parameter | Value | Description |
| --- | --- | --- |
| $p$ | $2 \times 10^{-7}$ cm/sec | The cell membrane's permeability to nitrocefin [41]. |
| $x_{am}$ | 132 cm<sup>2</sup>/mg | A conversion ratio equal to the cell's surface area divided by its mass [41]. |
| $k_{cat,p}$ | 10 s<sup>-1</sup> | Rate constant for nitrocefin export [41]. |
| $K_{M,p}$ | $4.95 \times 10^{-3}$ mM | Michaelis constant for nitrocefin export [41]. |
| $x_p$ | 0.3 | Fraction of the cell's volume that is consumed by the periplasm [34]. |
| $x_{th}$ | varies | The maximum concentration of nitrocefin in the periplasm that the cell can survive. |
| $k_{cat,E}$ | $4 \times 10^2$ s<sup>-1</sup> | Rate constant for nitrocefin hydrolysis [45]. |
| $K_{M,E}$ | $5 \times 10^2$ μm | Michaelis constant for nitrocefin hydrolysis [45]. |
| $sim\_data$ | see code | Parameters for wcEcoli. |

## Acknowledgments

This work was supported by the Paul G. Allen Frontiers Group via the Allen Discovery Center at Stanford and NIGMS of the National Institutes of Health under award number F32GM137464 to E.A. The content is solely the responsibility of the authors and does not necessarily represent the official views of the National Institutes of Health.

